# No evidence that human GIGYF2 interacts with growth factor receptor-bound protein 10 (GRB10): implication for human disease

**DOI:** 10.1101/2025.02.28.640929

**Authors:** Jung-Hyun Choi, Israel Shpilman, Niaz Mahmood, Shaghayegh Farhangmehr, Ulrich Braunschweig, Jun Luo, Reese Jalal Ladak, Angelos Pistofidis, T. Martin Schmeing, Seyed Mehdi Jafarnejad, Benjamin J. Blencowe, Nahum Sonenberg

## Abstract

GIGYF2 (Growth factor receptor bound protein 10 (GRB10)-interacting GYF (glycine-tyrosine-phenylalanine) protein 2) reduces mRNA stability and translation via microRNAs, ribosome quality control, and several RNA-binding proteins. GIGYF2 was first identified in mouse cell lines as an interacting partner with Growth factor receptor-bound protein 10 (GRB10), which binds to the insulin receptor and insulin-like growth-factor receptor 1. Mutations in the human *GIGYF2* gene were reported in autism spectrum disorder and Parkinson’s disease. In mouse models, mutations in the gene encoding GIGYF2 exhibited disease phenotypes. It was, therefore, thought that the GIGYF2-associated disease in humans is caused by defective GRB10 signaling. We show here that GIGYF2 does not interact with GRB10 in human cell lines, as determined by proximity ligation and co-immunoprecipitation assays. Size exclusion chromatography assay further confirmed these findings. The lack of interaction is explained by the absence of the critical GYF-domain binding PPGΦ sequence in the human GRB10 protein. These results are in contrast with the current understanding that a GIGYF2/GRB10 complex is associated with human disease via IR and IGF-1R signaling and underscore alternative mechanisms responsible for the observed phenotypes related to mutations in the human *GIGYF2* gene.

## Introduction

Mouse growth factor receptor-bound protein 10 (GRB10)-interacting GYF (glycine-tyrosine-phenylalanine) protein 2 (GIGYF2) was first identified as a GRB10-interacting protein through a yeast two-hybrid assay^1^. The gene encoding the orthologous human GIGYF2 protein is located on chromosome 2q37 and harbors CAG trinucleotide repeats. The human *GIGYF2* gene has a high pLI (probability of loss-of-function intolerance) score of 1, indicating robust genetic stability^2^. Mutations in the human *GIGYF2* gene are genetically linked to autism spectrum disorder (ASD). The Simons Foundation Autism Research Initiative (SFARI) gene database^3^ classifies mutations of *GIGYF2* as a category 1 (high confidence) risk factor for ASD (some of the mutations are shown in Fig. 1)^4-8^.

**Figure 1:**
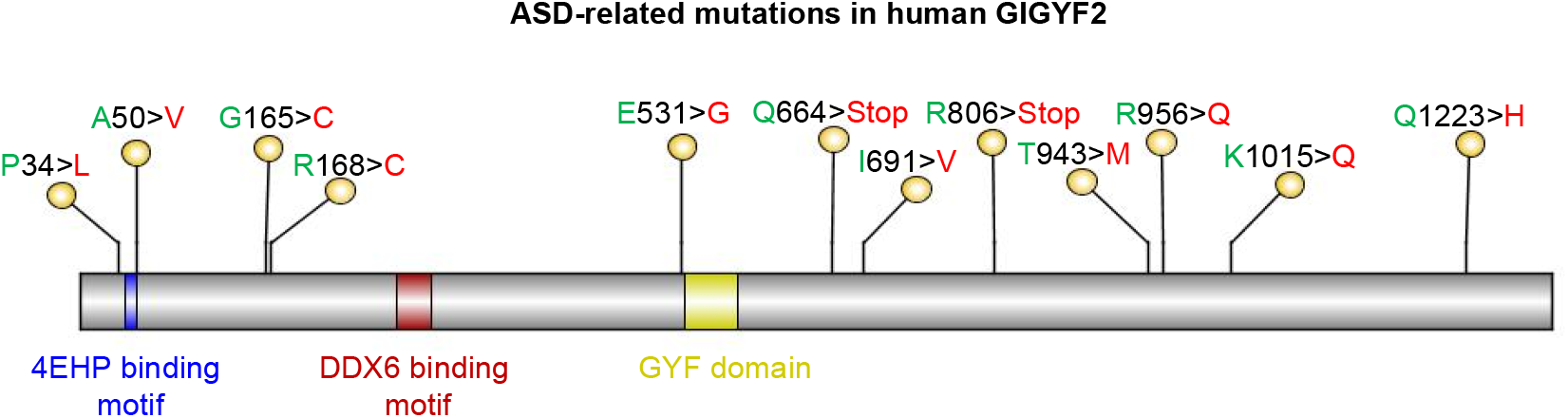
Key GIGYF2 mutations linked to autism. The different domains of GIGYF2 are illustrated as follows: DDX6 refers to DEAD (Asp-Glu-Ala-Asp) Box Helicase 6; GYF stands for glycine-tyrosine-phenylalanine domain.

GIGYF2 harbors several protein-binding motifs: the 4EHP (eIF4E homologous protein, or eIF4E2)-binding motif, DDX6 (DEAD-box helicase 6)-binding motif, and a GYF motif. GIGYF2 interacts with 4EHP via a conserved N-terminal motif (YXYXXXXLΦ, where Φ is a hydrophobic amino acid) to repress the translation of selective mRNAs in different biological contexts, including mouse embryonic development, activation of innate immune response upon viral infections, and upon binding of microRNAs (miRNAs) and RNA-binding proteins to mRNAs ^9-13^. The GYF domain interacts with partner proteins via the PPGΦ motif, where Φ is a hydrophobic residue (F/I/L/M/V)^14^. The PPGΦ containing GIGYF2-interacting proteins function in a variety of cellular processes via the trinucleotide repeat-containing 6 proteins (TNRC6A, TNRC6B, and TNRC6C) subunits of the microRNA-induced silencing complex (miRISC), the ribosome quality control factor zinc-finger protein 598 (ZNF598), and the RNA-binding protein tristetraprolin (TTP)^10-12,15,16^.

GRB10, a member of the GRB7 family of signaling adapters^17^, inhibits the phosphorylated tyrosine kinases bound to the insulin receptor (IR) and insulin-like growth factor 1 receptor (IGF-1R)^18^. This inhibition is linked to cognitive impairments associated with diabetes in mouse models ^19,20^. GRB10 in most mouse species (Muridae) contains a PPGΦ motif, which enables its interaction with GIGYF2^14^. It was reported that mouse GIGYF2 potentiates the GRB10-mediated inhibition of IR and IGF-1R function^1,20^. Mutations in the human *GRB10* gene are linked to ASD and the SFARI database^3^ designates it as a category 3 (suggestive evidence) risk factor. Strikingly, however, the PPGΦ motif is absent in human GRB10^14^. In this report, we show that, contrary to its designation as a GRB10-interacting protein, the human GIGYF2 does not bind to GRB10.

## Results

### The PPGΦ motif in GIGYF2 exists only in several species of the *Muridae* family

We examined GRB10 protein sequence alignments across vertebrates for the presence of the PPGΦ motif. We noted that the PPGΦ motif and its conserved flanking region are present only in several species of the *Muridae* family (Fig. 2). To investigate the cause for the absence of the PPGΦ motif in humans, we inspected the splicing pattern of orthologous *GRB10* genes in *Mus musculus* (house mouse; hereafter referred to as mouse) versus *Homo sapiens* (human). The PPGΦ motif in the mouse is encoded in an alternative spliced exon (transcript variant 1 and 3 containing exons 5 and 6, respectively, Fig. 3A). In sharp contrast, the corresponding sequence in orthologous human *GRB10* cannot serve as a coding exon. This is because both the donor and acceptor splice sites in the corresponding human sequence have diverged from the consensus splicing signals (5’ splice site GU changed to GC, and 3’ spice site AG diverged to GG, Fig. 3B). Most importantly, the human sequence orthologous to the murine exon contains a premature termination codon (in red, Fig. 3B). To validate that the latter orthologous sequence is not included in the mature mRNA, we examined the splicing patterns of *GRB10* transcripts in mouse versus human. Reverse transcription polymerase chain reaction (RT-PCR) was performed on RNA extracted from mouse brain cortex and human HAP1 (a near-haploid leukemia cell line) and RPE1 (retinal pigment epithelial-1) cells, using a pair of primers designed to amplify the corresponding human and mouse loci (Fig. 3C). Next, cycloheximide (CHX) treatment was used to inhibit the nonsense-mediated decay (NMD) pathway to enable the detection of potentially spliced but unstable or transiently expressed transcripts. We found that the exon encoding the PPGΦ motif is predominantly included in the mouse brain cortex (442 base pairs), but not in the human cell lines (Fig. 3D). As positive controls for CHX treatment and detection of NMD-degraded isoforms, RT-PCR analysis of premature termination codons (PTC)-containing isoforms transcribed from the serine/arginine-rich splicing factor 2 (*SRSF2*) and *SRSF6* genes was performed (Fig. S1). These data indicate that the PPGΦ motif is present in the protein encoded by mouse GRB10 but absent from the orthologous protein in humans. This raised the critical question whether GIGYF2 binds to GRB10 in humans.

**Figure 2:**
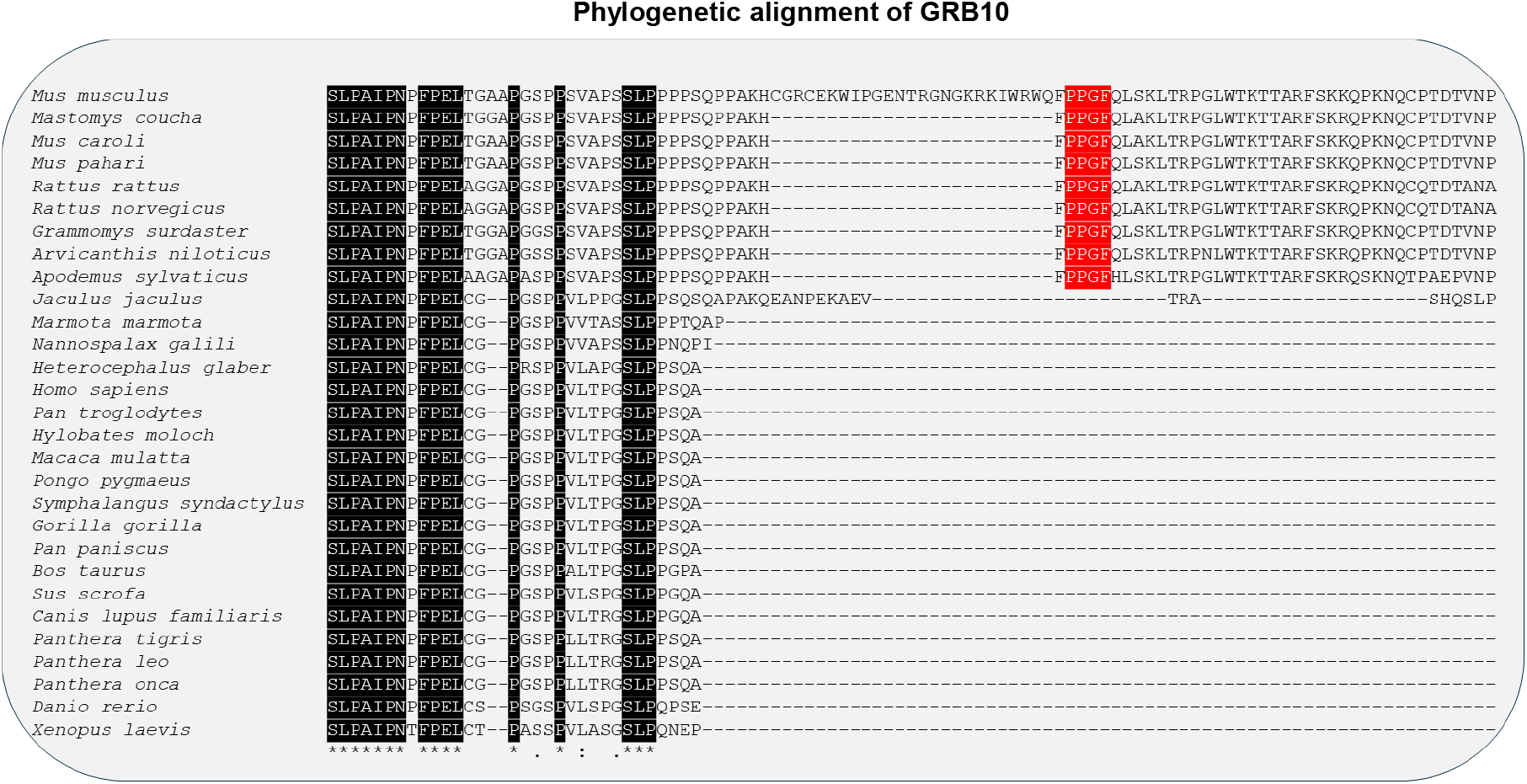
The PPGΦ motif of the GRB10 protein is present only in species of the Muridae family of rodents. Multiple sequence alignment (MSA) of GRB10 proteins of different species. The alignment was performed using Clustal Omega^35^. Amino acids that are similar across all analyzed species are shaded in black. The PPG motifs are highlighted in red. Due to space limitations, the complete protein alignment is not displayed here. The species names of the respective amino acid sequences are listed on the left. Amino acid sequences of GRB10 from a total of 29 species were analyzed. The binomial nomenclatures and common names (shown within brackets) are as follows: *Mus musculus* (House mouse), *Mastomys coucha* (Southern multimammate mouse), *Mus caroli* (Ryukyu mouse), *Mus pahari* (Gairdner’s shrewmouse), *Rattus rattus* (Black rat), *Rattus norvegicus* (Brown rat), *Grammomys surdaster* (African woodland thicket rats), *Arvicanthis niloticus* (African grass rat), *Apodemus sylvaticus* (Wood mouse), *Jaculus Jaculus* (Lesser Egyptian jerboa), *Marmota marmota* (Alpine marmot), *Nannospalax galili* (Middle East blind mole-rat), *Heterocephalus glaber* (Naked mole-rat), *Homo sapiens* (Human), *Pan troglodytes* (Chimpanzee), *Hylobates moloch* (Silvery gibbon), *Macaca mulatta* (Indochinese rhesus macaque), *Pongo pygmaeus* (Bornean orangutan), *Symphalangus syndactylus* (Siamang), *Gorilla gorilla* (Western gorilla), *Pan paniscus* (Bonobo), *Bos taurus* (Cattle), *Sus scrofa* (Wild boar/swine), *Canis lupus familiaris* (Dog), *Panthera tigris* (Tiger), *Panthera leo* (Lion), Panthera onca (Jaguar), *Danio rerio* (Zebrafish) and *Xenopus laevis* (African clawed frog).

**Figure 3:**
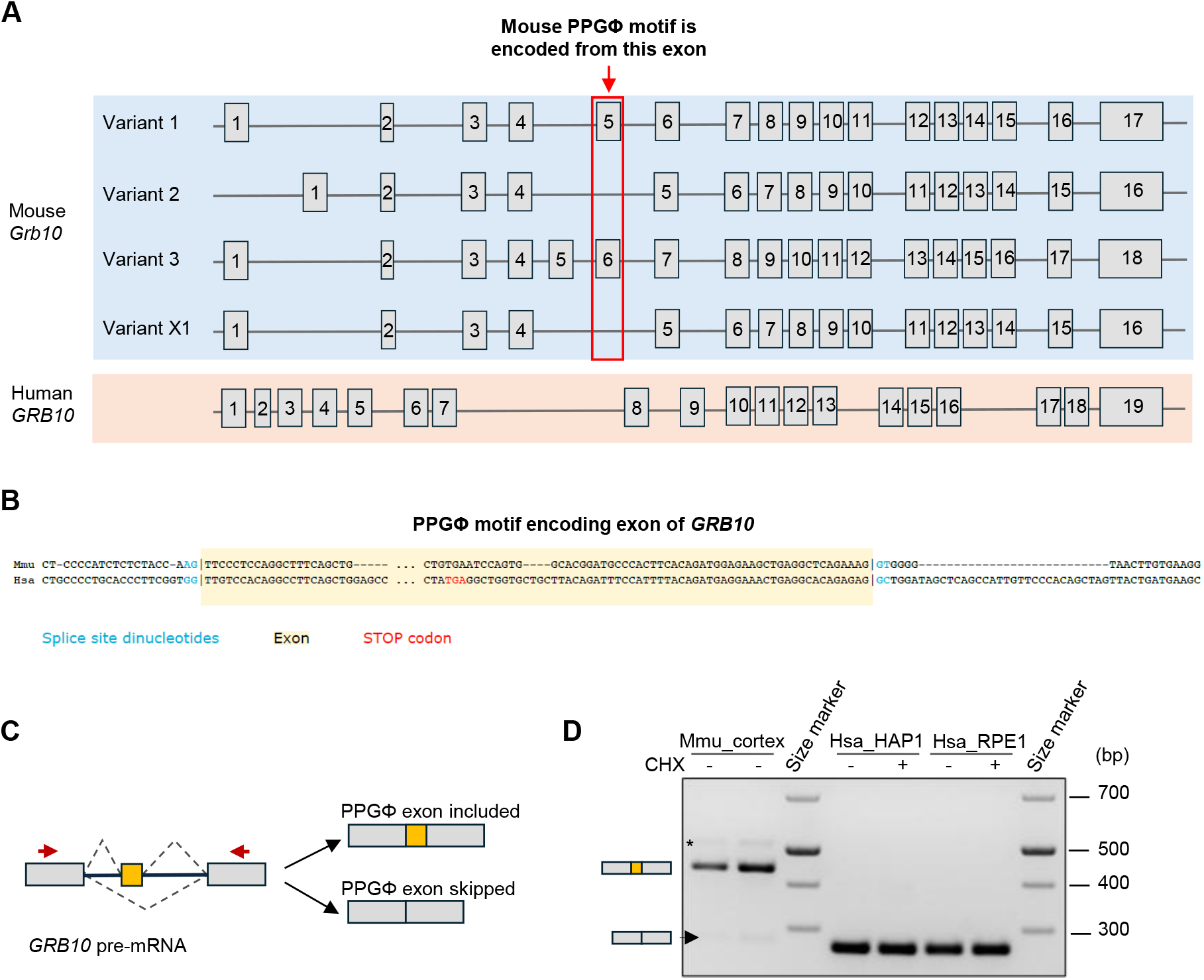
The PPGΦ motif of the human GRB10 is not present due to an alternative splicing event. (**A**) *Top:* Visualization of the *Grb10* exon-intron structure for mouse transcript variants obtained from Integrative Genomics Viewer (IGV) browser (mm10 genome), drawn from 5’ to 3’. Exons are depicted by grey boxes and are not drawn to scale. Mouse mRNA transcript variant 1 (NM_010345) and 3 (NM_001370603) contain exons 5 and 6 (respectively) by which the PPGΦ motif is encoded. Transcript variants 2 (NM_001177629) and X1 (XM_011243664) lack the exon which encodes the PPGΦ motif. *Bottom:* Visualization of the human *GRB10* exon-intron structure, drawn from 5’ to 3’. The human *GRB10* does not encode the PPGΦ motif. The human *GRB10* transcript variant 1 (NM_001350814) is shown (note that there are several other human transcript variants of *GRB10* present in the human genome, which are not shown here). **(B**) Comparison of the PPGΦ motif encoding sequences in human (Hsa) and mouse (Mmu) genes encoding GRB10. The splice sites are indicated in blue, and the alternative exon is marked by a yellow box shade. **(C**) The schematic model shows the design of a set of primers amplifying sequences of human and mouse *GRB10* genes flanking the exon encoding the PPGΦ motif. The exons and introns are represented by rectangles and lines, respectively. The exon encoding the PPGΦ motif is shown as a ‘yellow’ rectangle, while the flanking exons are colored in grey. Arrowheads depict the location of the forward and reverse primers used to detect the exon encoding PPGΦ motif. **(D**) RT-PCR analysis of total RNAs derived from mouse cortex and CHX-treated human HAP1 and RPE1 cells using primers described in **C**. The PCR products were resolved on a 2.5% agarose gel. The size of the skipped exon is 265 base pairs in humans and 277 base pairs in mice. The arrowhead indicates the skipped exon (minor band) in mice. Asterisk represents a band of unknown identity that is not the expected size for the target splice variant, which is only amplified in the samples of mouse origin.

### Absence of detectable interactions between human GRB10 and GIGYF2 proteins

We next assessed the interaction between human GRB10 and GIGYF2 by performing co-immunoprecipitation (co-IP) experiments in human embryonic kidney 293T (HEK293T) cells. All co-IP assays were conducted in the presence of ribonuclease A to eliminate potential RNA-bridging artifacts. While, as expected^12^, 4EHP and ZNF598 proteins co-precipitated with human GIGYF2, we failed to detect any co-precipitation of GRB10 and GIGYF2 (Fig. 4A). Conversely, IGF-1R co-precipitated with human GRB10 as expected^21^, but no co-precipitation of the GIGYF2 protein (Fig. 4B).

**Figure 4:**
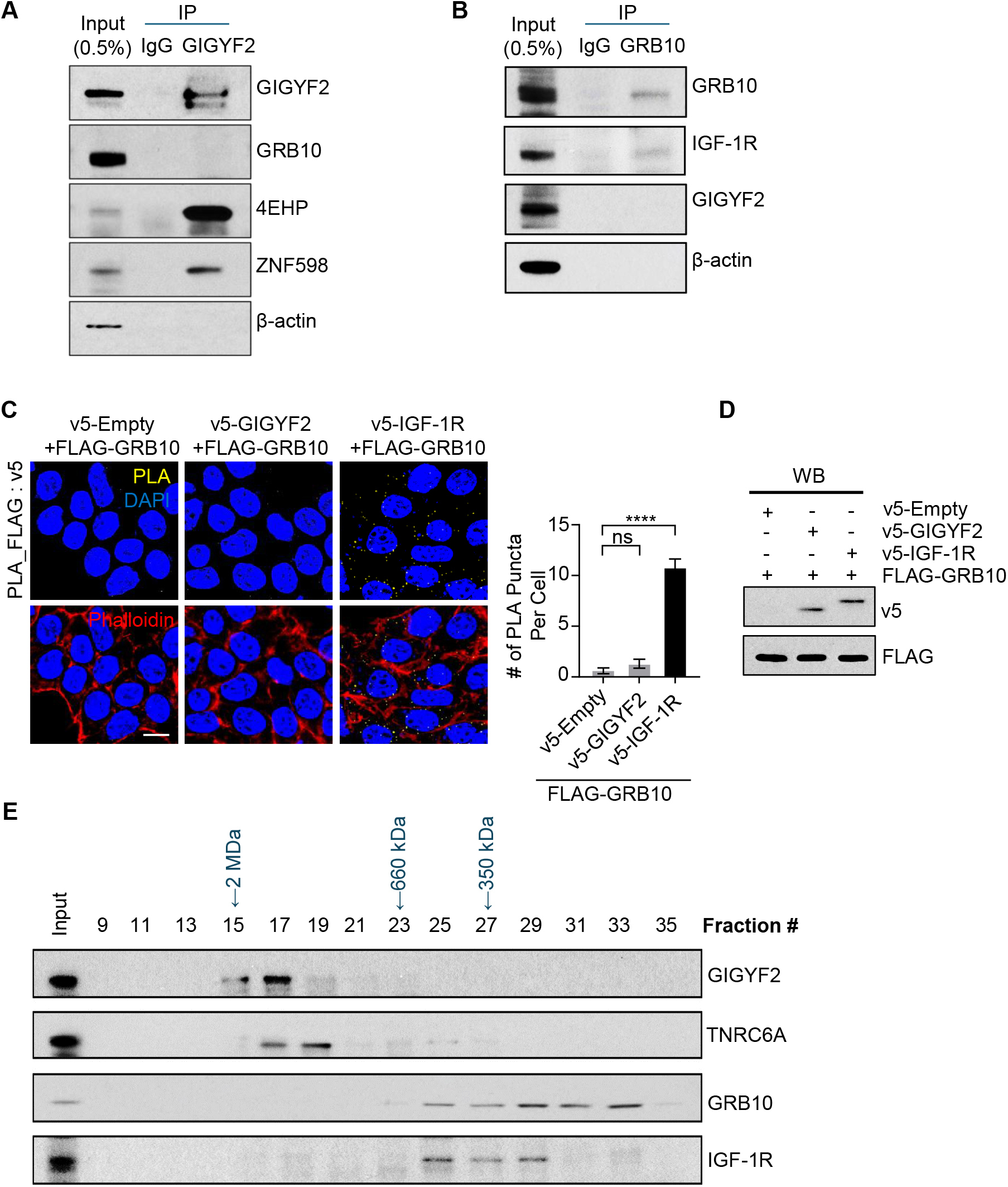
Analysis of interactions between human GIGYF2 and GRB10 proteins. (**A**) Co-immunoprecipitation (co-IP) analysis of endogenous GIGYF2 protein interactions in HEK293T cells. Immunoprecipitation was performed using an anti-GIGYF2 antibody, followed by western blot analysis with the indicated antibodies. (**B**) co-IP analysis of endogenous GRB10 protein interactions in HEK293T cells. Immunoprecipitation was performed using an anti-GRB10 antibody, followed by western blot analysis with the indicated antibodies. (**C**) Proximity ligation assay (PLA) for detecting interactions between FLAG-GRB10 and v5-IGF-1R or v5-GIGYF2. PLA signals are shown in yellow; the nucleus and actin cytoskeleton were counterstained with DAPI and phalloidin, respectively. Scale bar= 10 μm. The bar graph represents the average number of PLA signals from at least 30 cells per sample (n= 3 independent experiments). (**D)** Western blot analysis of cell lysates from **C**. (**E**) Analysis of endogenous GIGYF2 and GRB10 containing complexes by size-exclusion chromatography. A total of 10 mg of protein from HEK293T cells was loaded onto a Superose 6 column and ran at a flow rate of 0.4 mL/min. Fractions of 0.5 mL were collected, and 50 µL of each fraction was analyzed by western blotting.

To exclude the possibility that the lack of co-precipitation of human GIGYF2 and GRB10 proteins was due to technical limitations or unforeseen artifacts of the co-IP assay, we used an orthogonal validation approach with a proximity ligation assay (PLA). Corroborating the co-IP results, the co-transfection of FLAG-GRB10 with v5-IGF-1R yielded robust PLA signals (Fig. 4C & D). In contrast, minimal or no signals were detected in cells co-transfected with FLAG-GRB10 and either v5-Empty or v5-GIGYF2 (Fig. 4C & D). Taken together, these results indicate a lack of interaction between GRB10 and GIGYF2 in human cells.

### Human GIGYF2 and GRB10 proteins are components of distinct complexes

We next asked whether the human GRB10 and GIGYF2 fractionate in common complexes by examining the distribution of GIGYF2 and GRB10 proteins in native complexes via size exclusion chromatography (Superose 6) on lysates from human HEK293T cells. Our data revealed that human GIGYF2 and GRB10 proteins elute in distinct fractions (Fig. 4E). GIGYF2 primarily eluted in fractions associated with very high molecular mass complexes (>660 kDa) co-eluting with its known interactor TNRC6A (also known as GW182), a component of the miRISC (microRNA induced silencing complex)^15^. In sharp contrast, GRB10 and IGF-1R predominantly co-eluted in fractions corresponding to lower molecular mass (< 660 kDa). These differences in elution patterns demonstrate that, in the tested conditions, GIGYF2 and GRB10 are components of distinct protein complexes, suggesting divergent molecular functions.

AlphaFold3 generates high-quality models of protein complexes^22^. We submitted the protein sequences of human GRB10 and GIGYF2 to the AlphaFold3 server. As expected for noninteracting proteins, the five co-complex models^22^ had no commonality in the suggested GRB10: GIGYF2 interfaces, and each had very poor chain-pair interface-predicted template modelling scores (0.17 – 0.21) and minimum chain-pair-predicted aligned error scores (23.4 - 25.5). This indicates that AlphaFold3 does not predict the interaction between GRB10 and GIGYF2.

## Discussion

Our findings demonstrate that human GRB10 does not bind GIGYF2. This is non-surprising outcome considering the absence of the PPGΦ motif, which mediates the binding of the mouse GRB10 to GIGYF2 via its GYF motif. The exon encoding the PPGΦ motif of GRB10 is absent in humans, but it is predominantly included in Muridae due to changes in donor and acceptor splice sites in the latter during evolution. The Dependency Map (DepMap) portal of the Broad institute^23^ states that GIGYF2 might be involved in tyrosine kinase signaling. The STRING database^24^, a comprehensive resource providing information on known and predicted interactions, describes GIGYF2 as a GRB10-binding protein (Fig. S2). Considering our results, the entries where GIGYF2 and GRB10 proteins are mentioned as interacting partners must be amended

Studies on mouse models explored cognitive impairment due to diabetes through the repression of IGF-1R by GIGYF2^20^. GIGYF2 was shown to cause insulin resistance in obese mice by impairing the phosphoinositide 3-kinase/protein kinase B (PI3K/AKT) pathway^25^, which facilitates the translocation of glucose transporter GLUT4 from the cytoplasm to the cell membrane^26-29^. GRB10 inhibits IGF-1R and IR function independent of GIGYF2^19,30,31^. Knockdown of mouse *Grb10* augments downstream signaling pathways through the IGF-1R mediated phosphorylation of IR substrates, ERK1/2 and AKT^31^.

Because in humans, GRB10 cannot bind to GIGYF2, it is most probable that GIGYF2 and GRB10 function in humans through disparate pathways to inhibit the activity of IR and IGF-1R. Given these insights, further research on the relationship between GIGYF2 and GRB10 is necessary to clarify their role in insulin signaling and related pathologies. We recently described a mouse model in which 4EHP, the binding partner of GIGYF2, was depleted^32^. The mice exhibited a canonical ASD-like phenotype, including exaggerated hippocampal metabotropic glutamate receptor-dependent long-term depression (mGluR-LTD) and social behavior deficits^36^. In another study^15^, we have shown that the interaction between GIGYF2 and 4EHP is necessary for their mutual co-stabilization, as the loss of each protein results in the degradation of the other without affecting their mRNA levels. Thus, the cause of ASD in humans could be explained by the dysregulation of miRNA-induced translational silencing mediated by 4EHP due to changes in GIGYF2 instead of a putative GIGYF2/GRB10 axis. Thus, our findings are consistent with the involvement of 4EHP/GIGYF2 in the etiology of autism in humans.

## Materials and Methods

### Cell line and cell culture

HEK293T (Thermo Fisher Scientific) cells were cultured in Dulbecco’s Modified Eagle Medium (DMEM) (Wisent Inc., 319-005-CL) supplemented with 10% fetal bovine serum (FBS, R&D Systems, S12450) and 1% Penicillin/Streptomycin (Wisent Technologies, 450-200-EL). RPE1 cells were cultured in Dulbecco’s Modified Eagle Medium (DMEM) (Sigma-Aldrich, D5796) supplemented with 10% FBS (Gibco, 12483-020), 1 mM sodium Pyruvate (Gibco, 11360-070), and 1% Penicillin/Streptomycin (Gibco, 15140122). HAP1 cells were cultured in Iscove’s Modified Dulbecco’s Medium (Gibco, 12440053) supplemented with 10% FBS (Gibco, 12483-020) and 1% Penicillin/Streptomycin (Gibco, 15140122). To inhibit translation, cells were incubated in media either supplemented with 50µM cycloheximide solubilized in DMSO (Sigma-Aldrich, 01810) or DMSO for 6 hours prior to lysis. Cells were incubated in a humidified atmosphere of 5% CO2 at 37°C.

### Antibodies and plasmids

The following antibodies were used in the indicated dilution index: rabbit anti-GIGYF2 (1:3000, Bethyl, A303-732A-M), rabbit anti-GW182 (1:1000, Bethyl, A302-329A), rabbit anti-GRB10 (1:1000, Cell signaling, 3702), rabbit anti-IGF-1R (1:1000, Abcam, ab182408), rabbit anti-eIF4E2 (1:3000, Genetex, GTX82524), rabbit anti-ZNF598 (1:1000, Genetex, GTX119245), mouse anti-β-actin (1:5000, Sigma, A5441), mouse anti-v5 (1:3000, Abcam, ab27671), mouse anti-FLAG (1:5000, Abcam, ab49763), rabbit anti-FLAG (1:2500, Sigma, F7425). The λN-v5 tagged-GIGYF2 vector was described before^33^. v5 tagged-IGF-1R (Addgene, #98344) and FLAG-GRB10 (Addgene, #37481) were used for the PLA assay.

### RT-PCR

Total RNA was extracted from RPE1 and HAP1 cells using RNeasy Mini Kit (Qiagen, 74106) following the manufacturer’s instructions and including a DNase digestion step (Qiagen, 79254). Briefly, cells were washed twice with PBS and directly lysed in RLT buffer supplemented with 2-mercaptoethanol (Sigma-Aldrich, M3148). Embryonic mouse cortices were harvested at E18.5, lysed in RLT buffer supplemented with 2-mercaptoethanol, homogenized with QIAshredder (Qiagen, 79656), and subjected to RNA extraction as above. RT-PCR was performed according to the manufacturer’s instructions using a Qiagen one-step RT-PCR kit (Qiagen, 210212) using 10ng of total RNA for human cells and 20ng of total RNA for mouse cortices. The primer sequences used for RT-PCR are listed in Table S1.

### Western blotting

HEK293T cells were lysed with radioimmunoprecipitation assay (RIPA) lysis buffer (ThermoFisher Scientific, 89901), supplemented with a complete EDTA-free protease inhibitor (Roche, 04693124001) and phosphatase inhibitor (Sigma, P5726, P0044) cocktails. Protein quantification was performed using the Bio-Rad Protein Assay Dye Reagent Concentrate (Cat# 500-0006). Equal volumes of extracts were mixed with 2X-Laemmli sample buffer (Bio-Rad, Cat# 1610747), which included 5% β-mercaptoethanol. Proteins were denatured by heating at 95°C for 3 minutes and loaded onto a sodium dodecyl sulfate-10% polyacrylamide gel (SDS-PAGE) for electrophoresis. Proteins were transferred onto a nitrocellulose membrane, which was subsequently blocked using 5% milk in Tris-buffered saline with 0.1% Tween 20 (TBST, pH 7.6). Primary antibodies were applied to the membrane for 16 hours. The membranes were washed three times with TBST and incubated with horseradish peroxidase-conjugated secondary antibodies (Bio-Rad) for 1 hour. Signals were visualized using enhanced chemiluminescence (plus-ECL, PerkinElmer, Inc.) against an X-ray film (Denville Scientific, Inc.).

### Size exclusion chromatography

HEK293T cells were harvested by centrifugation and resuspended in 500 μL of lysis buffer containing 50 mM Tris-HCl (pH 7.4), 150 mM NaCl, 1 mM EDTA (pH 8.0), and 1% Triton X-100. The lysis buffer was supplemented with a complete EDTA-free protease inhibitor (Roche, 04693124001) and phosphatase inhibitor cocktails (Sigma, P5726, P0044). A Superose 6 Increase HR 10/300 column (Cytiva) was pre-equilibrated with SEC (size exclusion chromatography) buffer [25 mM HEPES-KOH (pH 7.5), 150 mM NaCl, 0.5 mM tris(2-carboxyethyl)phosphine [TCEP] at a flow rate of 0.4 mL/min. A total of 10 mg of protein from the HEK293T cell lysate was loaded onto the column. Fractions of 0.5 mL were collected, with 50 μL from each fraction used for western blot analysis.

### Immunoprecipitation

Cells were washed with cold PBS and collected by scraping into lysis solution (40 mM HEPES pH 7.5, 120 mM NaCl, 1 mM EDTA, 50 mM NaF, 0.3% CHAPS), supplemented with complete EDTA-free protease inhibitor (Roche, 04693124001) and phosphatase inhibitor (Sigma, P5726, P0044) cocktails. Pre-cleared lysates containing 2 mg of protein were incubated overnight at 4°C with 5 µg of anti-GIGYF2 or anti-GRB10 antibody, along with 40 µl of pre-washed Protein G agarose beads slurry (Millipore, 16-266) and RNase A (Thermo Scientific, EN0531). After the incubation, the beads were washed three times for 10 min each using a wash buffer (50 mM HEPES-KOH, pH 7.5, 150 mM NaCl, 1 mM EDTA, 50 mM sodium fluoride (NaF), 0.3% CHAPS), supplemented with complete EDTA-free protease inhibitor and phosphatase inhibitor cocktails. Proteins were eluted from the beads using the Laemmli sample buffer.

### Proximity Ligation Assay (PLA)

Proximity ligation assay (PLA) was performed using Duolink reagents (Sigma, DUO92101) following the manufacturer’s guidelines. Initially, cells were fixed in 4% paraformaldehyde (PFA) in sucrose for 15 minutes, followed by permeabilization with phosphate-buffered saline (PBS) containing 0.1% Triton X-100 for another 15 minutes. After permeabilization, the cells were blocked with Duolink blocking solution for 1 hour at 37°C and then incubated overnight at 4°C with primary antibodies. Following this, cells were washed with wash buffer A and treated with PLA probes for 1 hour at 37 °C. A ligation reaction was then carried out for 30 min at 37°C. The PLA signals were subsequently amplified using an amplification buffer for 100 min at 37°C. Finally, after a wash with wash buffer B, the samples were mounted onto glass slides and examined using an Airyscan microscope (Zeiss).

### Co-complex model generation

Prediction of co-complexes was performed with the AlphaFold3 via the AlphaFold Server (Beta) (https://alphafoldserver.com), using default seed auto-generation, which provides a set of 5 models^22^. Co-complexes modelled are of NP_001337743.1 (GRB10) and NP_001096617.1 (GIGYF2), and resulting model (.cif) and statstisiv (.json) files analyzed with ChimeraX^34^ and PyMOL (The PyMOL Molecular Graphics System, Version 3.0 Schrödinger, LLC).

## Statistical analysis

Statistical analysis was conducted using Prism 10 (GraphPad Prism Software Inc., USA). Error bars represent the standard deviation (SD) from the means of independent replicates. The number of replicates is specified in the figure legend. Statistical significance was set at p<0.05.

## Author contributions

JHC, IS, NM and SF performed experiments and data analysis. UB, JL, RL and AP assisted with the experiments and data analysis. TMS performed the AlphaFold3 analysis. TMS, SMJ, BJB and NS contributed to the design of the experiments. NS supervised the project. JHC, NM, IS, and NS wrote the manuscript. All authors edited and approved the manuscript.

## Acknowledgements

This work was supported by a grant from the Canadian Institutes of Health Research (CIHR; 148423) to NS and CIHR Project Grant 178084 to TMS. JHC was supported by a Conrad F. Harrington Fellowship. N.M. is supported by a postdoctoral fellowship from the Charlotte and Leo Karassik Foundation. The authors thank John Burke for AF3 tips. NS and TMS are members of the Centre de recherche en biologie structurale, funded by FRQS Research Centres Grant #288558.

## Figure Legends

**Figure S1: Detection of NMD isoforms in CHX-treated cells.** RT-PCR for *SRSF2* and *SRSF6* was performed as a positive control for the CHX assay, which inhibits the NMD pathway and allows for the detection of unstable or transiently expressed transcripts. The sizes of the bands corresponding to NMD isoforms are indicated in bold.

**Figure S2: The molecular interaction network of GIGYF2 protein predicted by the STRING database (https://string-db.org/).**

**Table S1:**
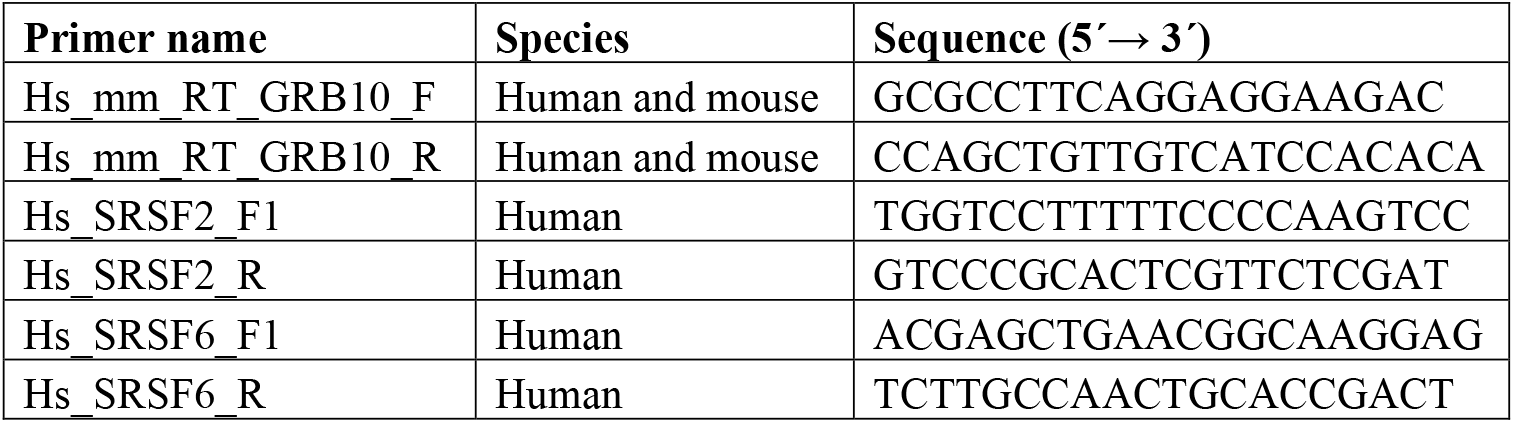
Primers used in the study are listed below:

