## Supplementary figures and images for "No evidence that human GIGYF2 interacts with growth factor receptor-bound protein 10 (GRB10): implication for human disease"

### Fig. S1

#### Supplementary Figure 1

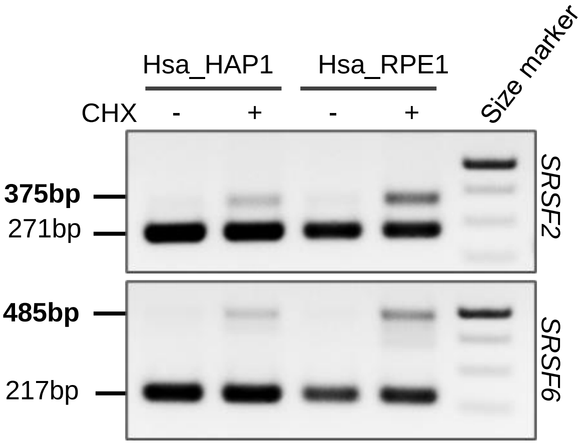

### Fig. S2

**Supplementary Figure 2**

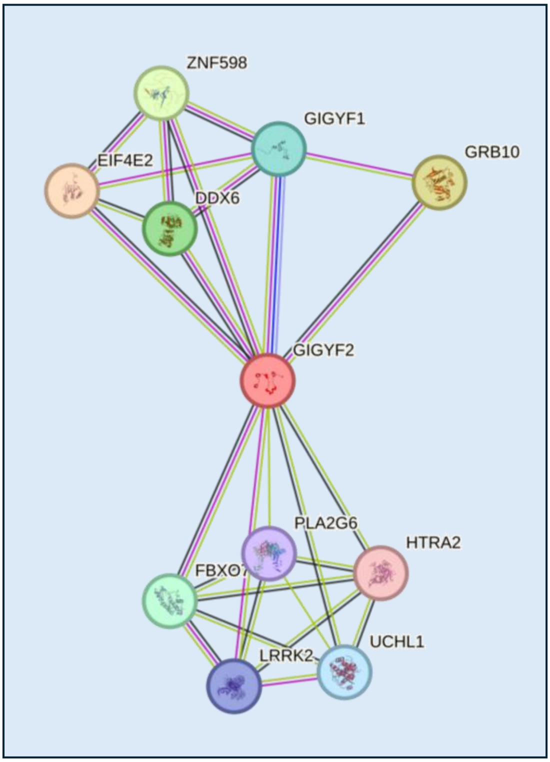
